# Minimally verbal children with autism may ‘see the point, but do not (always) point to what they see’: A behavioral and eye-tracking study in visual perceptual processing

**DOI:** 10.1101/2025.06.26.661808

**Authors:** Hannah S. Sykes-Haas, Yoram S. Bonneh

## Abstract

During typical development, non-social visual object recognition emerges in the first year of life, engaging low-level visual cues and higher-level mechanisms involving inference and prior knowledge. How these processes function in minimally verbal autism (mvASD) remains poorly understood. We studied children with mvASD (n=22, 6–11 years) using touchscreen-based oddball and contour-detection tasks targeting low-level stimuli (e.g. shape and orientation), and mid-level stimuli (e.g. illusory Kanizsa contours and 3D shapes). Pointing and eye-gaze responses were measured. Typically developing children (n=22, 6-12 years) served as a reference group. Accuracy and reaction-time profiles among mvASD participants were heterogeneous across experimental visual tasks and standardized developmental measures. All mvASD participants detected targets in the easiest condition, and approximately half succeeded across low-level tasks. Overall performance declined with increasing visual complexity, consistent with attenuated inference-based processing; communication ability and nonverbal reasoning together accounted for approximately 69% of between-participant variance in visual task performance. Critically, exploratory analyses suggested systematic perception–action dissociations rather than random error. First, the majority of participants who failed to point correctly (n=9) reliably fixated the correct target. Second, in the Kanizsa oddball task, nearly half of successful mvASD participants pointed to local inducers rather than the illusory figure center, unlike TDs. Third, more participants showed within-age-range nonverbal reasoning performance on Raven’s colorful progressive matrices when responding by puzzle placement than by pointing. These converging findings challenge interpretations of mvASD performance as reflecting perceptual or cognitive capacity alone, suggesting visual signals may guide action selection differently in mvASD.

**Lay Summary:** Minimally verbal children with autism showed individual differences in visual processing tasks. While developmental measures like communication ability and reasoning skills predicted most of the variation in performance, exploratory observations revealed an intriguing pattern: the same children sometimes succeeded when using their eyes to indicate answers but failed when pointing or performing better when placing puzzle pieces than pointing in a booklet to identical visual display. Several children who correctly detected illusory triangular shapes consistently touched the corner pieces rather than the triangle centers. These patterns suggest that performance depends not only on developmental and visual perceptual abilities, but also on how children are asked to respond. Parents and educators should consider: might a child who fails a pointing-based test succeed with a different response method?

## Introduction

> *AG, a beautiful six-year-old non-verbal girl with autism, still had no alternative augmentative communication (AAC) system in place. Despite intensive efforts, she struggled to find the image of a desired item even when only choosing between two. Inspired by a study in a “seemingly” unrelated field of visual crowding (Manassi et al., 2013), we modified the visual presentation of her cards by presenting the target card alongside seven blank cards. AG immediately selected the correct image, and she learned to discriminate desired items from distractor images as well. AG appears to have been challenged by visual <u>presentation and processing</u>, rather than semantic knowledge and the communicative nature of the AAC tool (first author HS, clinical case description)*.

During the first year of life, basic non-social visual object-recognition skills emerge in typically developing children (Haaf et al., 2003; Nishimura et al., 2009; McKyton et al., 2015). These abilities involve both low-level visual cues, such as color and orientation, and more advanced mechanisms relying on prior knowledge, integration, and inference, such as shape from shading (Bertone et al., 2005; Manassi et al., 2013; McKyton et al., 2015). In autism, these early visual mechanisms may diverge from typical developmental pathways (Gliga et al., 2015; Ostrolenk et al., 2017). Bertone et al. (2005) demonstrated superior perception of simple luminance-defined first-order gratings but diminished performance with more complex texture-defined second-order gratings requiring inference-based and integrative processing, leading to the ‘complexity-specific hypothesis’ of autism (Bertone et al., 2003, 2005). Similarly, Enhanced Perceptual Functioning (EPF) theory proposes that local low-level processing superiority may account for positive symptoms of autism, such as detail-focused perception, whereas reduced processing of complex perceptual material may contribute to negative symptoms, e.g. social-emotional reciprocity challenges (American Psychiatric Association, 2013; Mottron et al., 2006; Kéita et al., 2014). More broadly, extensive visual perceptual research in verbal autistic individuals supports superior local/simple processing, with some studies suggesting diminished global/complex processing (Frith and Happé, 1994; Dakin and Frith, 2005; Behrmann et al., 2006; Simmons et al., 2009; Booth and Happé, 2018).

Minimally verbal individuals with autism (mvASD), a subgroup representing the severe end of the autism spectrum and accounting for 25–30% of the autism population (Bal et al., 2016; Tager-Flusberg and Kasari, 2013; Rose et al., 2016), are treated extensively through visual aids, including visual AAC devices (Brignell et al., 2018), visual schedules, and picture exchange communication systems (PECS; Frost and Bondy, 2002; Arthur-Kelly et al., 2009; Hayes et al., 2010). Yet despite this ubiquitous reliance on visual support in clinical intervention, and despite the visual processing differences documented among verbal autistic individuals, how these perceptual patterns manifest in mvASD remain poorly understood.

Although there has recently been an upsurge in mvASD research (Courchesne et al., 2015; Bal et al., 2016; Brignell et al., 2018; Posar and Visconti, 2021; Girard et al., 2023; Pizzano et al., 2024; Kasari et al., 2025), very few studies have focused on visual perceptual processing within this clinical subgroup. To our knowledge, investigations into basic low-level visual processing among minimally verbal autistic individuals remain underexplored. Notably, Soulières and colleagues provide initial evidence that local visual strengths may also be present in mvASD, with typical or comparable performance on visual search and/or Children’s Embedded Figures tasks reported among some minimally verbal or largely minimally verbal autistic children (Courchesne et al., 2015; Girard et al., 2023). Given the complexity-specific hypothesis and EPF, and the suggested contribution of visual processing mechanisms to both positive and negative symptoms of autism, it is pertinent to examine whether differences in processing non-social, simple versus more complex low-level visual stimuli are also evident, or perhaps more pronounced, among the most severely affected clinical subgroup, namely children with mvASD. Investigating basic visual processing in mvASD is a critical starting point, as these mechanisms emerge early and provide a foundation for higher-order cognitive and perceptual functions.

Additionally, mvASD is increasingly recognized as heterogeneous in cognitive and behavioral profiles (Courchesne et al., 2015; Bal et al., 2016; Bauminger-Zviely et al., 2020; Pizzano et al., 2024), raising the question of whether visual processing patterns manifest uniformly or vary across individuals. Heterogeneity in mvASD has primarily been characterized through accuracy-based and categorical measures (Bal et al., 2016; Pizzano et al., 2024). Response dynamics, specifically, reaction-time (RT) sensitivity to visual task complexity, remain largely unexplored despite being a well-validated index of cognitive engagement in the typical population (Hick, 1952; Proctor and Schneider, 2018). Recent work in nonspeaking autism suggests that response timing can reveal systematic performance patterns (Jaswal et al., 2024).

In the current study, we investigated early visual processing in mvASD using touchscreen-based oddball and contour-detection tasks adapted from McKyton et al. (2015), with pointing on a touchscreen as the primary measure of target detection. This paradigm minimized verbal demands by allowing participants to imitate modelled task skills during practice, complete short, repeated trial runs, and respond with unlimited time. This approach is well suited for mvASD populations, with their complex and heterogeneous behavioral and cognitive profiles (Kasari et al., 2013; Tager-Flusberg et al., 2017; Matsuzaki et al., 2019; Bauminger-Zviely et al., 2020; Pizzano et al., 2024). In addition to accuracy, we examined RT across visual complexity levels as a candidate index of response-dynamics heterogeneity. During the course of the study, eye-gaze was incorporated as a complementary measure, motivated by concurrent work in our lab suggesting a possible dissociation between eye-gaze and pointing performance (Ellert et al., 2025). These analyses were exploratory.

We therefore asked: first, how do mvASD children perform across low- and mid-level visual tasks of increasing complexity? Second, how heterogeneous are accuracy and RT profiles across individuals? Third, to what extent is task performance associated with broader developmental measures, including communication ability, nonverbal reasoning and adaptive measures? Finally, in exploratory analyses, we examined whether eye-gaze and pointing responses reveal perception–action dissociations that may inform interpretation of task performance in mvASD. Findings indicate that mvASD children are not a homogeneous group, and that performance may not solely reflect broad developmental and visual perceptual ability, but rather how visual information is translated into action, suggesting a dissociation between perception and action.

## Methods

### Participants

#### Experimental group

Twenty-eight children with mvASD who met inclusion criteria (see below) were initially recruited. Six were excluded after testing began due to safety-related behavior (2), motor limitations preventing reliable pointing responses (n=2), uncorrected vision (refusal to wear prescribed glasses, n=1), and verbal output during testing was inconsistent with the caregiver-reported mvASD classification (n=1). The final sample comprised 22 participants (16 males, 6 females; age range 6.9–11.1 years, M = 8.9 ± 1.3 years). The age range reflects both developmental and practical considerations. The task demands, such as sustained cooperation with a touchscreen and eye-tracker across multiple sessions, require behavioral readiness that children with mvASD typically acquire only after several years in the special education system (Tager-Flusberg et al., 2017).

The children were recruited from three special-education schools and pre-schools in Israel with a relatively large cohort of students with mvASD. All mvASD participants met inclusion criteria for the autism group. Each child had received a formal clinical ASD diagnosis according to DSM-5 criteria (American Psychiatric Association, 2013), supported by standardized diagnostic information from the ADOS and/or an ADI-R–based developmental interview (Lord et al., 1994, 2000). Diagnosis was given by two independent accredited autism diagnosticians, as required by Israeli Health Ministry and Ministry of Education for entry into the special education school system for students with autism. At least one diagnostician was a physician (child neurologist, psychiatrist, or developmental pediatrician), and the second was either a physician or developmental/clinical child psychologist. None were affiliated with the study. The ASD diagnosis was further verified through teacher-reported scores above the autism cutoff (≥15) on the Social Communication Questionnaire (SCQ; Rutter et al., 2003).

#### Raven’s Matrices Colored Progressive Matrices

MvASD participants underwent assessments of fluent reasoning using Raven’s Colored Progressive Matrices in both booklet and puzzle board form (RCPM; Raven et al., 1998). The RCPM is believed to be a core measure of fluid intelligence, assessing abilities such as rule inference, goal management, and abstract reasoning independent of language ability (Dawson et al., 2007; see Table 1 for summary of descriptive measures and appendix 1 for descriptive measures for individual participants). For three mvASD participants (Pseudonyms Levi, Lia, and Haim), RCPM performance data is missing due to leaving the school or behavioral challenges (see Appendix 1). These cases were excluded from analyses involving RCPM but retained for all other analyses where applicable. Missing data were not imputed.

**Table 1.** Summary of descriptive developmental measures^1^.

| mvASD<br>(n=22) | Mean (SD)<br>(excluding f.p.<sup>2</sup>.) | Range | Estimated IQ |
| --- | --- | --- | --- |
| Boys/Girls | 16/6 | ---- |  |
| Chronological age in months | 107 (16) | [83;133] |  |
| L-VIS (communicative competence)<sup>3</sup> | 8.4 (3) | [2;14] |  |
| RCPM puzzle<sup>4</sup> (n=9)-<br>nonverbal fluid reasoning-percentiles | 50 (35) | [f.p.;97<sup>th</sup>] | [f.p.;129] |
| RCPM Booklet<sup>4</sup> (n=5)-<br>nonverbal fluid reasoning- percentiles | 66 (36) | [f.p.;97<sup>th</sup>] | [f.p.;128] |
| Perseveration on the RCPMpuzzle<sup>5</sup><br>(same 2 consecutive answers) | 7.5 (6.5) | [0;20] |  |
| SCQ score<sup>6</sup> | 24 (4) | [15;30] |  |
| Vineland-3<sup>7</sup>: |  |  |  |
| - Communication | 40.8 (9.2) | [25;58] |  |
| - Daily Living Skills | 37.7 (11.8) | [20;60] |  |
| - Socialization | 31 (10.9) | [20;59] |  |
| - Motor Skills | 54.6 (11.8) | [36;74] |  |
| - Adaptive Behavior Composite (ABC) | 20.4 (1.5) | [20;27] |  |

<sup>1</sup> RCPM data not available for some mvASD participants (see Appendix 1 for more details)
<sup>2</sup> f.p.=floor performance
<sup>3</sup> Maximum possible score on L-VIS communicative abilities is 17.
<sup>4</sup> Mean calculated excluding f.p. (less than 1<sup>st</sup> percentile) and only scores within age-range (above 5<sup>th</sup> percentile. No mvASD scored between 1<sup>st</sup> and 5<sup>th</sup> ).
<sup>5</sup> All participants with a RCPMpuzzle f.p. made at least 3 mistakes by perseveration (see appendix 1).
<sup>6</sup> Cut-off score for increased likelihood of autism is $\geq 15$ (Rutter et al., 2003).
<sup>7</sup> Vineland-3 Domain-Level Teacher Form standard scores (normative mean=100, SD=15); lower scores indicate greater adaptive impairment.

#### Low-Verbal Investigatory Screener

Parents completed the Low-Verbal Investigatory Survey 4.0 (L-VIS 4.0; Naples et al., 2023, see also: https://autismlanguagelab.psychology.uconn.edu/low-verbal-investigatory-survey-elvis-project-home), a newly developed brief (5-minute) caregiver-report tool assessing communication in minimally verbal autistic children. In this study, L-VIS 4.0 was used solely to assess communication abilities. Although the original factor analysis supports a two-factor model, only the first factor, communication abilities, was statistically validated and analyzed. This factor reflects typical verbal and receptive communication, with high internal consistency and strong correlations with standardized measures (Vineland-II: 0.73; PLS-5: 0.61; Stanford-Binet non-verbal: 0.53). The second factor, atypical communication, was not used for composite scoring due to mixed loadings and interpretive concerns. However, we report one item from this factor, production of unusual sounds (e.g., shrieks, as it was a strong predictor of performance across multiple gold-standard assessments (PLS, VABS, and Stanford-Binet IQ). For three participants (See appendix 1) the L-VIS 4.0 was completed by trained care staff familiar with the child for over a year, due to lack of parental response.

#### Vineland Adaptive Behavior Scales, Third Edition (Vineland-3)

Adaptive functioning was assessed via the Vineland-3 Domain-Level Teacher Form (Sparrow et al., 2016), with teachers who had worked with each participant for at least one year, rating them across Communication, Daily Living Skills, Socialization, and Motor Skills domains.

#### mvASD criterion

Participants met the criterion for minimal spoken language if caregivers (parent and teachers) indicated that they either rarely or never used meaningful phrase-level speech spontaneously in everyday situations, or if their functional spoken vocabulary consisted of roughly 30 or fewer distinct communicative words or phrases, not including echoed language (Plesa Skwerer et al., 2016). According to the L-VIS, twenty participants were defined by parents or familiar staff as non- or minimally verbal, 14 as having ‘no spoken words’ and 7 as having ‘some spoken words’. One participant (‘Hans’) was described as having ‘many spoken words’, but these were echolalic and he had less than 30 spontaneously spoken communicative words (see Appendix1).

#### Exclusion criteria

Participants were excluded if they had known comorbid developmental, neurological, or genetic conditions, with the exception of medically controlled epilepsy. Reports of developmental delay and intellectual disability was not criterion for exclusion, as standardized tests and/or diagnostician’s clinical impression may at times underestimate cognitive abilities in mvASD (Courchesne et al., 2015, Bauminger-Zviely et al. 2020, Pizzano et al 2024).

Four of the 22 remaining participants did not complete all experimental tasks. One male mvASD participant was excluded from two oddball tasks because of emerging safety concerns, while a female mvASD participant moved abroad before finishing the final task. Two other participants joined the study later and only participated in selected tasks.

Sixteen children in the autism group were receiving psychotropic medications to manage behavioral challenges, maintaining focus and/or sleep regulation. They were prescribed Risperidone (Risperdal), Medical Cannabis, Carbamazepine (Tegretol) Ritalin, Atent and melatonin. All mvASD participants were assigned pseudonyms for easier identification across tasks (see Results).

#### Reference group

A reference group of twenty-two typically developing (TD) children participated in the study (age range: 6-12 years; M = 9.3 ± 2.1 years), comprising 12 males and 10 females. Only 14 were tested on the Kanizsa oddball condition, which was added later. None of the reference group participants had any known developmental, neurological, cognitive, or genetic conditions, and all had normal or corrected-to-normal vision according to parental report. Descriptive measures were not collected for TD participants, who served as a reference group. The visual abilities probed by these tasks are functionally present early in life (Gerhardstein & Rovee-Collier, 2002); ceiling-level performance was expected and confirmed (see Results), rendering additional descriptive measures uninformative.

Ethics approval was obtained from the Chief Scientist of the Israeli Ministry of Education and the Ethics Committee at Bar-Ilan University. Informed consent was obtained from all parents (mvASD and TD), and data collection took place at either the child’s home or at their educational institution per parental preference. All TDs and one mvASD participant were tested in their homes.

### Apparatus

Testing was conducted using a Lenovo Yoga C940 laptop featuring a 14-inch touchscreen display with a resolution of 1920 x 1080 pixels and a refresh rate of 60 Hz. Eye-tracking data was recorded using the Tobii Eye Tracker 4C, operating at a 90 Hz sampling rate and providing gaze tracking accuracy of 0.5-1 degrees. The Tobii eye-tracker was selected for its non-invasive nature, allowing participants freedom of movement while maintaining tracking accuracy, as demonstrated in our recent study with patients (Shani et al., 2025). The experimental setup was stabilized using a Tobii EyeMobile computer tray mounted on an adjustable monitor arm. This configuration ensured consistent positioning and protected the devices from sudden unpredictable behaviors. Stimulus presentation was managed through the PSY platform, an in-house software for psychophysics and eye tracking developed by Y.S.B. (second author).

### Stimuli

All tasks involved visual oddball or contour detection arrays presented on a touchscreen. The stimuli spanned a range of visual perceptual complexity, from basic low-level features (e.g., luminance contrast, color hue) to mid-level (e.g., 3D shading, Kanizsa figures, contour integration), similar to McKyton et al., (2015). Oddball stimuli were displayed in 2×3 and 2×2 grid configurations at fixed screen locations, with standardized viewing distance (~50–60 cm), visual angles (~2.5°–6.5°), and fixed inter-stimulus spacing. The stimuli are presented in Figure 1, showing the actual stimulus displays for the 2×3 configurations, with trials differing only in the position of the oddball. These displays were all ~10 cm (center-to-center) in width, except for the displays in Figure 1b1 (~17 cm), a4 and b4 (~13 cm). All the other sizes and positions were in proportion to the display size as shown. The backgrounds were white, gray, or black (luminance in cd/m^2^, a1: 1.5; a2-3: 75; a4, b4: 110; b1: 335; b2-3: 100). The patches in a3 were a white target, and black, dark grey, or mid grey distracters (145, 3, 31, 61 cd/m^2^ respectively). The grating stimuli in a4 were square patches of 1 cpd sine-wave gratings with 32% contrast, vertical with horizontal oddball. The stimuli in b4 were a second-order version of a4, created by a sine-wave modulation of the contrast of a random dot pattern (1-pixel texels with random luminance) as in Bertone et al. (2005).

**Figure 1.**
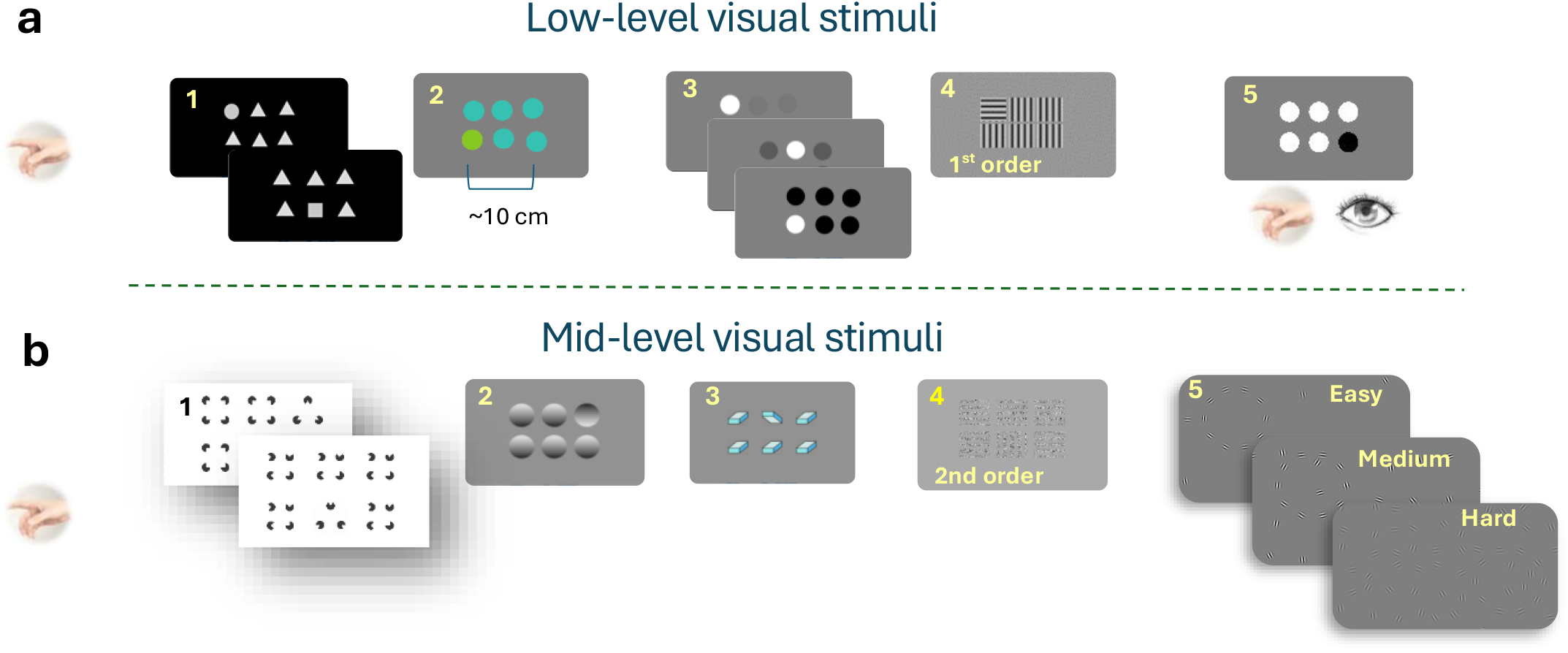
The experimental paradigms and sample stimuli. **(a)** Low-level oddball paradigms. **(a1)** shapes (**a2**) Color (**a3**) Black/white saliency (**a4**) 1^st^ order orientation gratings, (**a5**) white/black w/eye-gaze. **(b)** Mid-level oddball and contour paradigms., **(b1)** Kanizsa, **(b2)** Light from above, (**b3**) 3D box orientation, (**b4**) 2^nd^ order orientation gratings (**b5**) Circular contours with increasing distractor density.

The contour detection stimuli (Figure 1b5) were generated as configurations of symmetric Gabor patches (wavelength l and envelope s of 16 pixels) with 30% contrast on a gray (60 cd/m^2^) background, similar to Bonneh & Sagi (1998), but in which a circle is embedded in a random background (Field et al., 1993). First, a background of randomly oriented Gabor patches arranged on a grid with a specific spacing that varied across conditions was set to cover 1900×600 pixels rectangle, with a position jitter radius per patch of 48 pixels. Then, a circle of patches with a radius of 160 pixels, inter-patch interval of 88 pixels (5.5 l), and uniform tangential orientation jitter of +/-20 deg, was added in one of 4 positions (+/-450 and +/-250 pixels for x and y respectively). Background patches that were closer than 48 pixels to a target patch (center-to-center) were erased. There were 3 contour difficulty levels in separate runs according to the background inter-patch interval: Easy (640, 192 pixels), Medium (128, 96 pixels), and Hard (64,48 pixels), two intervals mixed per level. Full stimulus parameters are provided in Table S1 of the Supplementary material.

### Procedure

#### Experimental pointing tasks

Testing was conducted in a quiet room (home or school) with lighting at ~500 lux. The experimenter (first author, HSH), an experienced clinician in mvASD, ran all sessions. Participants sat ~60 cm from the screen and could move freely. The experimenter sat to their right, with rewards placed 70–80 cm away. Each participant completed 8–10 sessions over 3–8 months. Sessions lasted 10–15 minutes, with breaks based on child’s need and motivation. Participants could leave at any time. Sessions were scheduled no more than once weekly, with longer gaps due to holidays or absences. Before each block, participants received brief verbal instructions. For oddball tasks, they were told to “find/point to the one that’s different”; for the circular contour task, “find/point to the circle.”

There were 7 oddball pointing conditions, 3 circular contour detection by pointing conditions, and one oddball task requiring gaze *and/or* pointing (Figure 1a and 1b). To address language limitations, each condition began with 8–10 practice trials. The experimenter modelled pointing to the correct item and after 3–4 failed modelled practice trials she would use hand-over-hand guidance (if the child consented). Each condition had 10–32 trials, with oddball targets randomly positioned in 2×2 or 2×3 grids at any of the 4 or 6 possible locations. Conditions were kept brief (10–30 trials, ~20s–2min) to maintain engagement and allow breaks. Trial counts per run were: Color, shapes, BWG: 4 runs × 10 trials; First-/Second-order: 2 runs × 20 trials; Box, light-from-above: 4 runs × 18 trials; Kanizsa: 1–2 runs × 24 trials; Contour detection: 3 difficulty levels, 16 trials each (4 locations of contour × 2); tested once or twice per participant (48–96 trials total). Actual trial numbers varied due to performance, cooperation and trial rejection. In pointing tasks, new trials began upon screen touch. Trials occasionally included rapid repeated touches (double-clicks), which were handled during data preprocessing (see Data Analysis). During the experiments, rewards (e.g. food, toys, videos) were tailored based on caregiver input and provided for task success and cooperation.

#### Exploratory analyses

In addition to the planned analyses, we conducted exploratory analyses motivated by observations arising during data collection and by concurrent findings in our lab suggesting a possible dissociation between perception and action.

#### Eye-gaze vs. pointing

Eye-gaze assessment was added as an exploratory extension, motivated by a concurrent finding from our lab: minimally verbal autistic individuals demonstrated accurate written word recognition via eye-gaze while performing at chance when pointing to the same targets (Ellert et al., 2025). This gaze-pointing dissociation raised the question of whether a similar pattern might emerge in basic visual perceptual processing.

Eye-tracking was conducted in a subset of mvASD participants who performed at or near chance on multiple pointing tasks during the main experimental study (see Figure 2a-h). This procedure yielded a final eye-tracking sample of n = 9 participants (two additional low-performing participants could not be included due to relocation and safety constraints). These participants correspond to the low-performing subgroup identified in the data-driven analyses reported in the Results section. We combined simultaneous pointing and eye-gaze recording with an eye-gaze-only paradigm to minimize occlusion of the eye-tracking signal. Given time and participant constraints, a single condition was selected for this paradigm: a luminance-reversed oddball display relative to the original pointing task (black oddball among white distractors; see Figure 2d for the non-reversed display and corresponding performance), chosen based on their near-chance performance in the corresponding non-reversed condition. This luminance-reversed version was used to assess performance under a comparable task structure while avoiding repetition of identical stimulus displays. This high-salience, single-feature (“pop-out”) condition is typically resolved via preattentive parallel processing in typical populations (Treisman & Gelade, 1980; Wolfe, 1994). In the black/white eye-gaze oddball task, each run included 12 trials. Participants completed between 3 and 16 runs. Valid trial counts varied across participants (range: 7–75 trials), reflecting differences in cooperation and data quality. Some trials were unusable due to movement, looking away, or eye-tracker obstruction. Analysis was only conducted on eye-gaze data within the 2×3 stimulus display region. In the eye-gaze-only condition, displays remained visible for 2.5 seconds.

**Figure 2.**
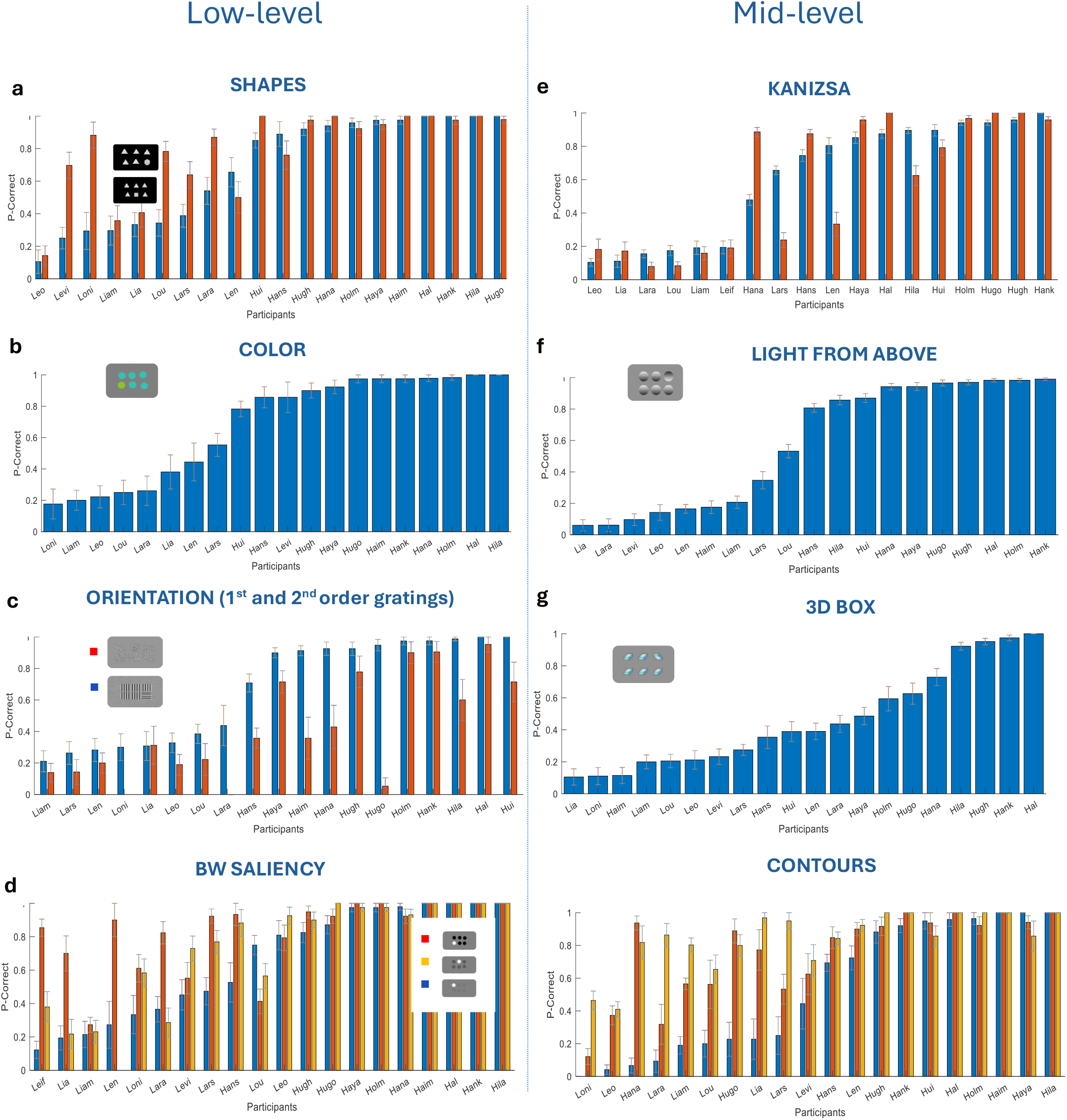
Performance on the oddball and contour detection tasks via pointing. **(a)** shapes, **(b)** color, **(c)** 1^st^ and 2^nd^ order orientation gratings **(d)** Black/white saliency, **(e)** Kanizsa, **(f)** light from above and **(g)** 3D box orientations and **(h)** circular contours among increasing no. of Gabor distractor elements (see Figure 1 for visual stimulus). Data were plotted with a bar for each mvASD participant sorted from low to high performance in one stimulus condition in each experiment (e.g. Shape experiment sorted according to square oddball condition). Pseudonyms were used for each participant (see results section ‘Heterogeneity’) with initials “L” for low and “H” for high performers. Error bars denote 1SE of the mean across trials.

Eye-tracking used the standard 3-point Tobii 4C calibration, supported by experimenter guidance or reward-based fixation cues. Calibration quality was immediately verified using the Tobii verification display, with a manual verification by the experimenter of within-circle fixation, implying accuracy better than ~1.5 deg. The percentage of valid gaze samples (excluding blinks and off-screen artifacts) was high across the cohort (M = 82%, SD = 11). Spatial precision, quantified as the Root Mean Square (RMS) of sample-to-sample (S2S) deviations during fixation, was calculated to assess instrumental noise. Median RMS sample-to-sample gaze noise was 0.16° (IQR: 0.046°). Eye-tracking data were not collected from the TD reference group or high-performing mvASD participants, as performance in these groups was at or near ceiling on this oddball task.

#### Kanizsa off-center pointing

During testing of the circular contour conditions, we observed anecdotally that a number of mvASD participants who successfully detected the target contour consistently pointed to the contour’s frame rather than its center (off-center pointing was not systematically recorded in the circular contour conditions). Given documented inclinations toward local processing in ASD even in the presence of intact global perception (Mottron et al., 2006), and consistent with this anecdotal pattern observed in the preceding circular contour condition, we predicted that some participants who successfully detected the global Kanizsa illusory triangle would similarly tend to point to local Pac-Man inducers rather than the globally defined figure’s center, unlike TDs, the majority of whom were expected to point centrally. To test this, we recorded pointing location relative to the triangle center (i.e., off-center pointing) for all participants who successfully detected the Kanizsa illusory triangle (individual mean > 0.5; chance ≈ 0.15).

### Data Analysis

All analyses were conducted at both the participant and trial levels, depending on the research question. Accuracy was defined as proportion correct (P-Correct), and analyses were restricted to valid trials as defined below.

#### Pointing

For oddball tasks, correct responses were defined as touches within a 150-pixel radius of the target center. For contour detection, correct responses were defined within a 200-pixel radius of the target center. To ensure validity, only touches occurring more than 200 ms after the previous touch were included, as some participants produced rapid repeated touches (“double-clicks”) in which the second touch landed on an incorrect location and did not reflect a genuine response.

#### Eye-gaze

Two complementary measures of eye-gaze accuracy were defined, applied to data recorded during the 2.5 s display window. (1) **Transient fixation accuracy (TFA)**: fixation within a 200-pixel radius of the target center during a 300–700 ms window following stimulus onset. (2) **Sustained fixation accuracy (SFA):** cumulative fixation time on the target exceeding that on any distractor within a 0.4–1.5 s window after stimulus onset, capturing competitive fixation dynamics. The choice of the time range was taken from the mvASD group mean RT during pointing, which was ~1.5 s. For both measures, trials were included only if gaze fell within a central 400 × 500 pixel region, which corresponds to the full stimulus array region.

The 200-pixel spatial criterion was adopted to accommodate reduced calibration precision and fixation stability due to participant movement, consistent with known gaze-tracking challenges in autism (e.g., Wass et al., 2015). While this relatively liberal threshold may reduce spatial specificity, it provides a consistent and conservative basis for comparing eye-gaze and pointing performance across modalities. As a sensitivity check, a wider 220-pixel criterion was also evaluated (see Results).

#### Reaction times

Reaction times (RTs) were recorded for all trials, measured from display onset. Accuracy analyses included all responses occurring more than 200 ms after display onset. RT analyses were restricted to correct trials with RTs below 2500 ms, to exclude trials with extended behavioral delays (e.g., children standing up or walking away before returning to respond) while retaining valid responses.

#### Descriptive developmental measures

<u>Nonverbal reasoning</u> was assessed using the RCPM in both booklet and puzzle board form. Raw scores were converted to age-normed percentiles (Raven et al., 1998), interpolated linearly, and scaled to approximate IQ scores (IQ = 100 + z×15), as in Courchesne et al. (2015). <u>Perseveration</u> on the RCPM puzzle was defined as selecting the same answer location on two consecutive items; the RCPM never places the correct answer in the same location on adjacent items, so such responses indicate position-based repetition rather than correct reasoning. <u>Communication ability</u> was assessed using the L-VIS 4.0 Communication Abilities factor (maximum score = 17; Naples et al., 2023). Autism symptomatology was indexed by the SCQ total score, with all participants scoring above the autism likelihood cutoff of ≥15 (Rutter et al., 2003). <u>Adaptive behavior</u> was assessed using the Vineland-3 Domain-Level Teacher Form (Sparrow et al., 2016). Raw domain scores (Communication, Daily Living Skills, Socialization, and Motor Skills) were converted to age-normed standard scores (normative mean = 100, SD = 15) using the Domain-Level Teacher Form norm tables (Appendix D; Sparrow et al., 2016). The Adaptive Behavior Composite (ABC) score was derived as the sum of the Communication, Daily Living Skills, and Socialization domain standard scores. Lower scores indicate greater adaptive impairment.

#### Heterogeneity: Subgrouping

Subgrouping was performed to characterize heterogeneity within the mvASD sample and examine whether performance patterns differed systematically between subgroups. Subgrouping was based on a single participant-level variable: mean pointing accuracy across all visual tasks. To characterize heterogeneity in performance, the distribution of participant-level mean accuracy was first assessed for normality (Shapiro–Wilk test) and unimodality (Hartigan’s dip test). Gaussian mixture models (GMMs) with one to three components were then fitted, and model fit was compared using Bayesian Information Criterion (BIC). The number of components was not specified a priori; the final solution was selected based on model fit and interpretability, with preference for a solution that did not yield trivially small subgroups.

#### Statistical analysis

Trial-level data were analyzed using linear mixed-effects models (LMEs), with random intercepts for participant and session nested within participant. Random slopes were included where supported by model convergence. Although correct/incorrect responses are binary at the trial level, LMEs were used to test overall patterns across repeated observations while accounting for within-participant dependence. To assess whether this modeling choice affected the main conclusions, we repeated the primary trial-level accuracy analyses using logistic generalized linear mixed-effects models (GLME) with a binomial distribution and logit link. The direction and significance pattern of the main effects were unchanged.

Depending on the analysis, fixed effects included stimulus level (low vs. mid), group (mvASD and TD), response modality (pointing vs. eye-gaze), and standardized descriptive measures (RCPM, L-VIS, SCQ, VABS-3). For categorical predictors, reference levels were specified so that low-level stimuli, TD participants, and pointing responses served as the reference categories unless otherwise noted. Thus, coefficients for stimulus level reflect the change from low-to mid-level stimuli, group coefficients reflect differences relative to TD participants, and modality coefficients reflect the gaze–pointing difference. Reaction times were analyzed using LMEs with fixed effects of group, stimulus level, and their interaction.

Eye-gaze and pointing performance were compared using paired-sample t-tests at the participant level and confirmed using trial-level LMEs including response modality as a fixed effect and random intercepts and slopes for participant. Off-center pointing in the Kanizsa task was analyzed at both the participant level (independent-samples t-test) and the trial level (LME with pointing distance from the triangle center as the dependent variable and group as a fixed effect), to confirm that results were consistent across levels of aggregation. The primary inferential tests were the mixed-effects models specified above; follow-up pairwise tests were treated as exploratory and interpreted with caution, with emphasis on effect sizes and convergence across analytic levels. Effect sizes (Cohen’s d) are reported for all t-tests. Statistical significance was evaluated at α = .05.

Standardized measures included communication ability (L-VIS), nonverbal reasoning (RCPM), autism symptom severity (SCQ), and adaptive functioning domains from the Vineland-3: Communication, Daily Living Skills, Socialization, and Motor Skills. Because these measures varied only at the participant level and several were conceptually related, we examined their interrelationships prior to model specification (Figure S1). This analysis showed substantial intercorrelations among L-VIS, RCPM, and Vineland adaptive-functioning domains, indicating overlapping variance across these measures, whereas SCQ showed little association with task performance or with other standardized measures. To avoid overparameterization and unstable estimates given the limited number of participants, the primary mixed-effects models focused on two theoretically motivated predictors: communication ability (L-VIS) and nonverbal reasoning (RCPM).

## Results

### Descriptive overview of task performance across visual levels

In the oddball and contour detection tasks participants were required to detect, via pointing, visual stimuli among distractors. Performance was measured as the proportion correct (P-Correct) for each mvASD participant across eight different visual stimuli. Figure 2 illustrates the individual performance accuracy (P-Correct) for mvASD in each condition, displayed in ascending order from lowest to highest performance. Each bar represents a single participant. In stimulus tasks with two or more conditions, ordering was sorted according to one of the conditions (e.g. shapes task according to performance on square oddball). The results show variability in accuracy among participants and visual levels, with some participants showing near chance level performance and others at or near ceiling accuracy rates.

To investigate performance across different levels of visual stimuli, we compared accuracy between low- and mid-level stimuli, categorized following McKyton et al. (2015). Trial-level accuracy was analyzed using a linear mixed-effects model (Table 2, Model 1). Performance was significantly lower for mid-level compared to low-level stimuli (β = −0.163, p < .0001), indicating reduced accuracy with increasing perceptual complexity (see Figure S2). Typically developing children (TDs) performed at or near ceiling (oddball P-Correct: 0.98–1; contour: 0.95–1, not shown).

**Table 2.** Summary of trial-level and participant-level statistical models. Trial-level models used linear mixed-effects models with random effects as specified in Table S2. Participant-level analyses were used as complementary between-subject tests. Marginal R^2^ reflects variance explained by fixed effects; conditional R^2^ includes random effects. **Notes:** (1) Model formulas and variable definitions are provided in Table S2. R^2^m = marginal R^2^; R^2^c = conditional R^2^, (2) for model 6, Group was coded such that the negative coefficient reflects greater off-center pointing in mvASD relative to TD.

| Model | Level | Outcome | Predictors | N | Key result | Model test / variance explained |
| --- | --- | --- | --- | --- | --- | --- |
| 1 | Trial | Accuracy | Stimulus level | 9,581 | Mid-level < low-level, $\beta = -0.163$ | $F(1,9579) = 186.63$, $p < .0001$ |
| 2 | Trial | Accuracy | Group × level | $\sim 9,581$ | No interaction, $p = .47$ | — |
| 3 | Trial | RT | Group × level | 8,670 | Level effect attenuated in mvASD, especially LowP | $F(5,8664) = 30.20$, $p < .0001$ |
| 4 | Trial | Accuracy, TFA | Modality | 218 | Gaze > pointing, $\beta = 0.283$ | $F(1,218) = 7.76$, $p = .006$ |
| 5 | Trial | Accuracy, SFA | Modality | 218 | Gaze > pointing trend, $\beta = 0.192$ | $F(1,218) = 3.10$, $p = .08$ |
| 6 | Trial | Off-center pointing | Group | 1,399 | mvASD more off-center than TD; $\beta = -42.02$ negative $\beta$ reflects coding direction | $F(1,1397) = 22.36$, $p < .0001$ |
| 7 | Trial | Accuracy | RCPM + L-VIS-C | 8,081 | Both positive predictors | $R^2_m = .19$; $R^2_c = .44$ |
| 8 | Subject | Mean accuracy | RCPM + L-VIS-C | 18 | L-VIS-C positive predictor, $\beta = 0.0506$, $p = .001$; RCPM borderline, $\beta = 0.0026$, $p = .050$; | $R^2 = .69$ |
| 9 | Subject | TFA gaze vs pointing | Modality | 9 | Gaze > pointing | $t(8) = 3.19$, $p = .013$, $d = 1.05$ |
| 10 | Subject | SFA gaze vs pointing | Modality | 9 | Gaze > pointing trend | $t(8) = 1.99$, $p = .081$, $d = 0.70$ |
| 11 | Subject | Off-center pointing | Group | 26 | mvASD < TD | $t(24) = -4.54$, $p < .001$, $d = 1.79$ |

### Individual differences, subgrouping and response dynamics

Overall performance across all visual tasks was non-normally distributed (Shapiro–Wilk W = 0.86, p = .006), and Hartigan’s dip test rejected unimodality (D = 0.116, p = .010). To characterize the structure of this heterogeneity, we compared Gaussian mixture models (GMM) with one through three components. A three-component solution provided a marginally better fit than two components (ΔBIC = 14.35 vs. 11.28, relative to a single Gaussian) but yielded one subgroup with very few participants (n = 2), limiting interpretability. The remaining participants were split into two larger groups comparable to the two-component solution. We therefore retained the two-component solution, which captured the main structure of the data. The two components corresponded to a low-performing group (LowP-mvASD, n = 11; M = 0.38, SD = 0.09; range 0.23–0.52) and a high-performing group (HighP-mvASD, n = 11; M = 0.89, SD = 0.08; range 0.68–0.99), with no participant scoring between 0.52 and 0.68 (Figure 3). Participants were assigned pseudonyms.

**Figure 3.**
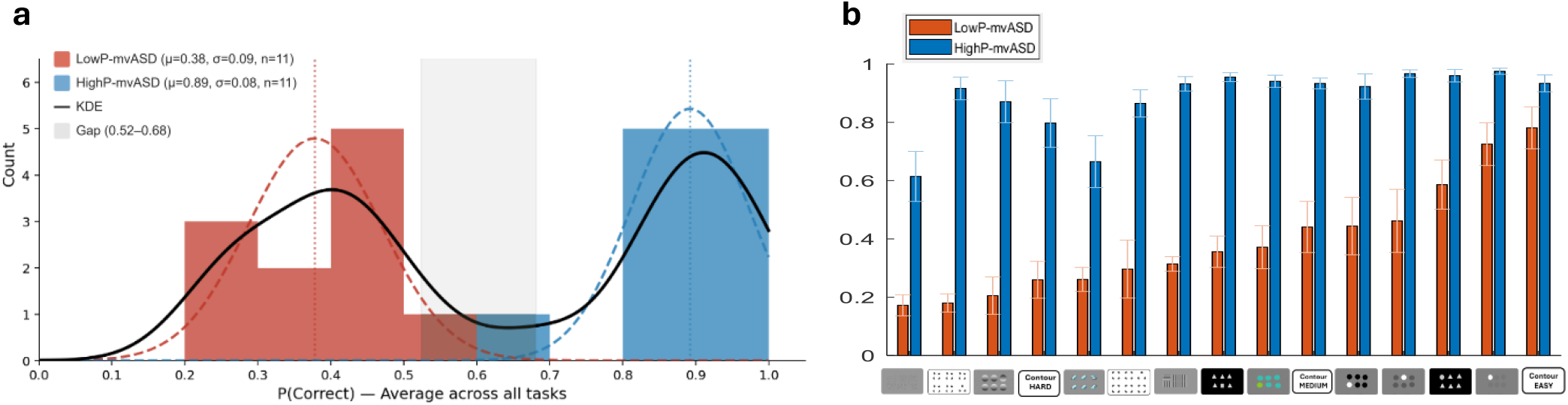
Distribution of overall visual task performance in children with mvASD suggests data-driven subgrouping. **(a)** Histogram of mean performance (P-Correct) averaged across all visual tasks for 22 mvASD participants. Red and blue bars indicate participants assigned to the low- and high-performing subgroups (LowP-mvASD and HighP-mvASD), respectively, based on Gaussian mixture modeling (see Results). Dashed curves show the two-component GMM fit; the solid black curve shows the kernel density estimate (KDE) of the full distribution. The shaded gray region marks the gap between 0.52 and 0.68, within which no participant scored. Dotted vertical lines indicate group means (low: μ = 0.38; high: μ = 0.89). **(b)** Mean P-Correct in all low and mid-level visual tasks among LowP-mvASD (red bars) and HighP-mvASD (blue bars). Because subgroup assignment was based on overall mean P-Correct, panel b is shown descriptively and was not treated as an independent test of group differences.

Following the data-driven identification of LowP- and HighP-mvASD subgroups, we tested whether the stimulus-level effect on trial-level accuracy differed between groups using a linear mixed-effects model. The group × level interaction was not significant (Table 2, Model 2; β = −0.022, p = .47; full model specification in Table S2), indicating that accuracy declined from low-to mid-level stimuli to a similar extent in the two mvASD subgroups.

Finally, we analyzed reaction times (RTs) on correct trials as a function of subgroup and visual task level. The trial-level LME showed a significant overall effect of the fixed factors (Table 2, S2, Model 3). Across tasks, LowP-mvASD participants showed consistently longer RTs than TD participants (β = 235.33 ms, p < .001), whereas HighP-mvASD did not differ from TD participants (β = 16.44 ms, p = .78), closely tracking the TD group (Figure 4a). RTs increased with task level in TD participants (β = 286.17 ms, p < .0001), indicating slower responses for mid-level visual tasks. This level-dependent RT increase was reduced modestly in HighP-mvASD (β = −95.79 ms, p = .02), but was strongly attenuated in LowP-mvASD (β = −270.02 ms, p < .0001). Thus, RT modulation by task level was largely preserved in HighP-mvASD but markedly reduced in LowP-mvASD. To visualize this interaction more directly, Figure 4b, shows RT as a function of task level averaged across tasks. As seen, TDs exhibited a marked increase in RT from low-level to mid-level. HighP-mvASD showed a more moderate increase, and LowP-mvASD showed little change across levels despite overall slower responses. This pattern is consistent with the reduced level-dependent modulation captured by the interaction terms in the model.

**Figure 4.**
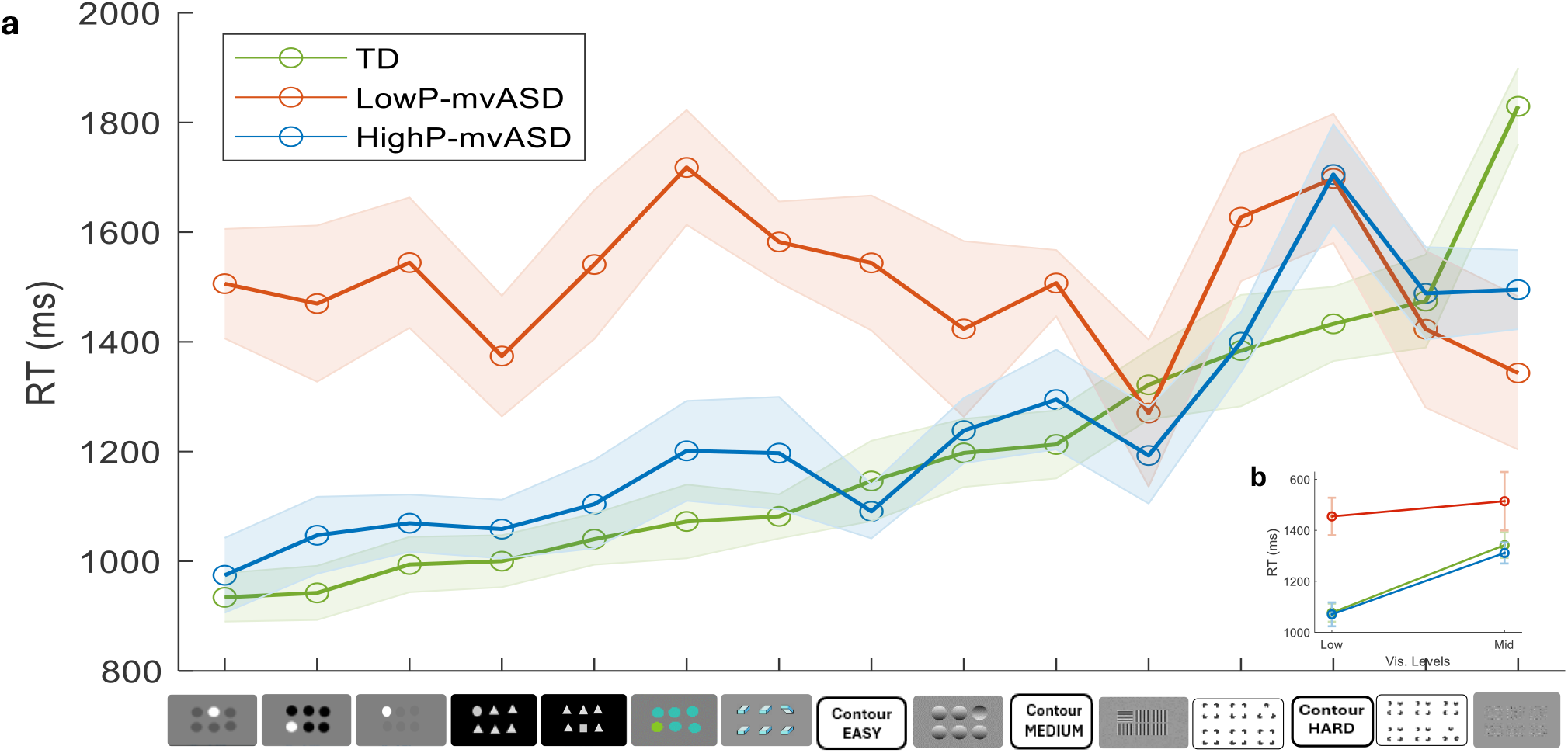
Reaction time across tasks and task levels in TDs, LowP-mvASD and HighP-mvASD: **(a)** Mean reaction time (RT) for each task (N=8670), plotted separately for TD reference group (green), HighP-mvASD (blue), and LowP-mvASD (red). Tasks are ordered according to increasing RT in the TD reference group. Error bars represent ±1 SE across participants. The LowP-mvASD show consistently longer RTs across tasks, whereas the HighP-mvASD closely follows the TDs. **(b)** Mean RT as a function of task level (low-level vs. mid-level), averaged across tasks. Error bars represent ±1 SE. TDs show a marked increase in RT with visual level, HighP-mvASD show a reduced increase, and LowP-mvASD show minimal modulation of RT by visual level despite overall slower responses. This pattern corresponds to the group × level interaction observed in the linear mixed-effects model.

### Exploratory analysis 1: Eye-gaze vs. pointing in lower performing mvASD subgroup (LowP-mvASD)

In an exploratory analysis, we compared eye-gaze and pointing performance in LowP-mvASD participants on the saliency oddball task (Figure 5). Using the transient fixation accuracy measure (TFA; see Analysis), eye-gaze performance was higher than pointing in both the participant-level paired comparison and the trial-level LME accounting for unequal trial counts across participants (Table 2, Models 4 and 9). The trial-level model showed a significant gaze advantage (β = 0.283, p = .006), and the participant-level comparison showed a similar effect (p = .013). The convergence of these analyses provides preliminary evidence for a gaze–pointing dissociation in the LowP-mvASD subgroup.

**Figure 5.**
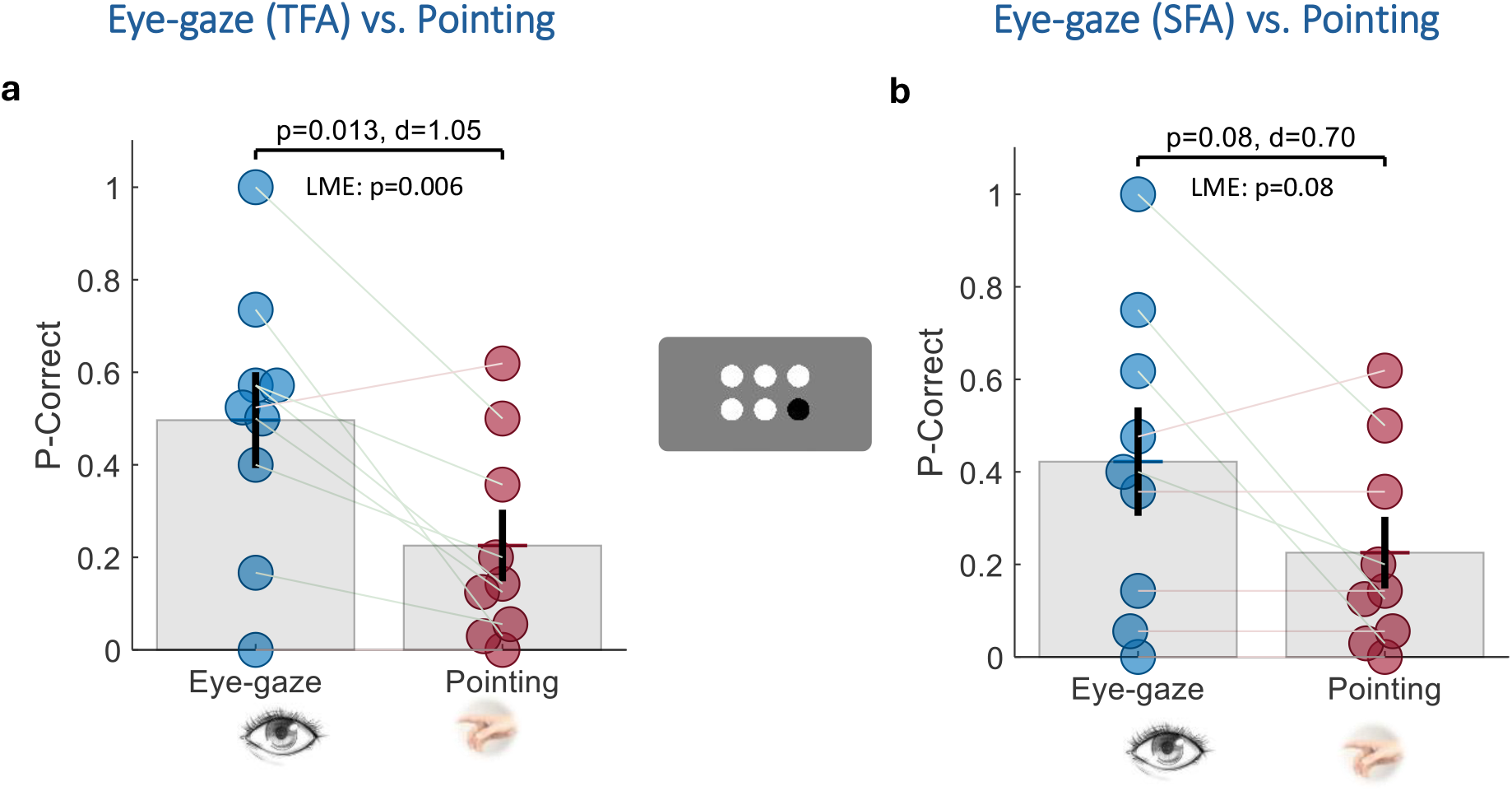
Eye-gaze vs. pointing in LowP-mvASD subgroup. Each pair of circles connected by a gray line represents an individual participant’s performance (n = 9) in the black/white saliency oddball task. Participants were instructed to “touch or look at the odd one out.” **(a)** Transient fixation accuracy (TFA), defined as fixation within a 200-pixel radius of the target center in the 300–700 ms window. **(b)** Sustained fixation accuracy (SFA), defined as cumulative fixation time on the target exceeding that on any distractor (0.4–1.5 s window). Eye-gaze performance was consistently higher than pointing, reaching significance under TFA and showing the same directional pattern under the stricter SFA criterion. Error bars represent ±1 SE. The p-value from the participant-level group comparison, the LME p-value, and Cohen’s d are shown above each panel.

Using the more conservative sustained fixation accuracy measure (SFA; see Analysis), eye-gaze performance showed the same directional advantage over pointing, but this effect did not reach significance in either the participant-level comparison or the corresponding trial-level LME (Table 2, Models 5 and 10). The trial-level model showed a non-significant gaze advantage (β = 0.192, p = .08), and the participant-level comparison showed a similar trend (p = .081). A sensitivity analysis using a wider spatial criterion (220 pixels) yielded stronger effects for both TFA and SFA, but this criterion was not retained because it overlapped with nearby distractor regions and reduced fixation specificity. Together, the two measures indicate a consistent gaze advantage in LowP-mvASD, reaching significance only under the less conservative TFA criterion.

### Exploratory Analysis 2: Atypical pointing in the Kanizsa oddball task

Consistent with the prediction derived from the contour-pointing observation, off-center correct pointing during detection of the triangular Kanizsa oddball was observed in both mvASD and TD participants. Notably, two participants from the LowP-mvASD group also performed well in this condition (Figure 2e). Off-center pointing distance differed significantly between groups in both the trial-level LME and the participant-level comparison (Table 2, Models 6 and 11). In the trial-level model, mvASD participants showed reduced central pointing relative to TDs (β = −42.02, p < .0001), with a consistent participant-level group difference (p < .001). TD participants consistently pointed near the center of the illusory triangle, whereas mvASD participants showed a more bimodal pattern, pointing either to the triangle center or to the centers of the Pac-Man inducers (Figure 6a,b).

**Figure 6.**
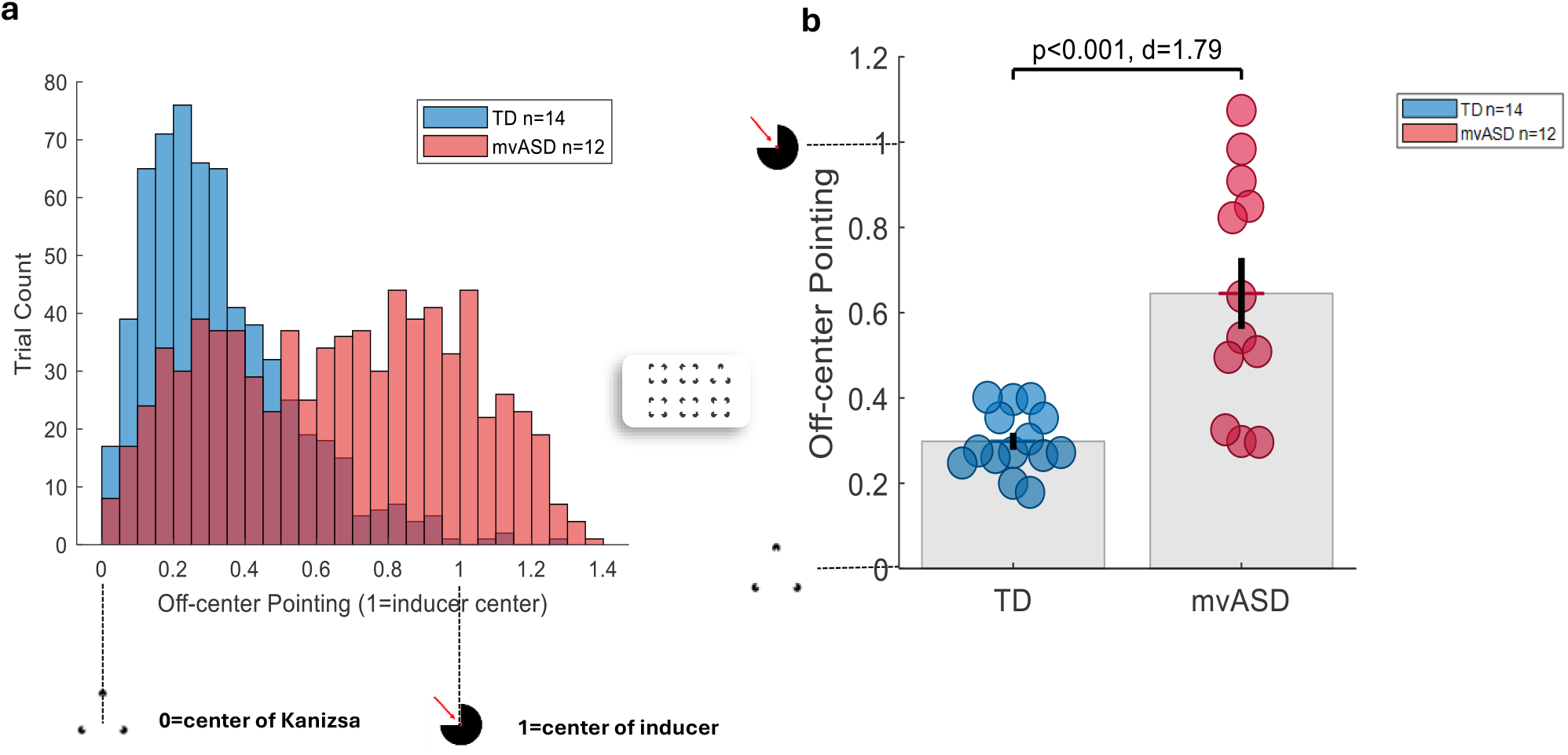
Off-center pointing on target Kanizsa oddball. Off-center pointing refers to pointing distance from the center of the illusory triangle, normalized to the range 0–1, where 0 = center of the Kanizsa triangle and 1 = center of a Pac-Man inducer; values exceeding 1.0 indicate pointing beyond the inducer center. Analysis included all TD participants tested on the Kanizsa condition (n = 14) and mvASD participants performing above chance (n = 12). **(a)** Trial-level histogram of off-center pointing. In contrast to TDs, mvASD pointing was distributed around the Kanizsa center <u>and</u> the inducer center. **(b)** Participant-level group comparison. Each circle represents one participant. Error bars represent ±1 SE. The p-value from the participant-level group comparison, the LME p-value, and Cohen’s d are shown above panel (b).

### Standardized descriptive developmental measures associations to task performance

We first examined descriptive bivariate associations between pointing-based visual task accuracy and standardized assessment scores using Spearman correlations (Figure 7; Figure S1). Accuracy was positively associated with L-VIS-C communication scores (ρ = 0.79, p = .0001) and RCPM Puzzle scores (ρ = 0.61, p = .016) and negatively associated with perseverative responses on the RCPM Puzzle task (ρ = −0.65, p = .005). SCQ scores were not associated with task accuracy (ρ = −0.11, p = .633). Among Vineland-3 domains, accuracy was positively associated with Daily Living Skills (ρ = 0.57, p = .016) and Motor Skills (ρ = 0.50, p = .032), but not with Communication (ρ = 0.44, p = .072) or Socialization (ρ = 0.19, p = .447).

**Figure 7.**
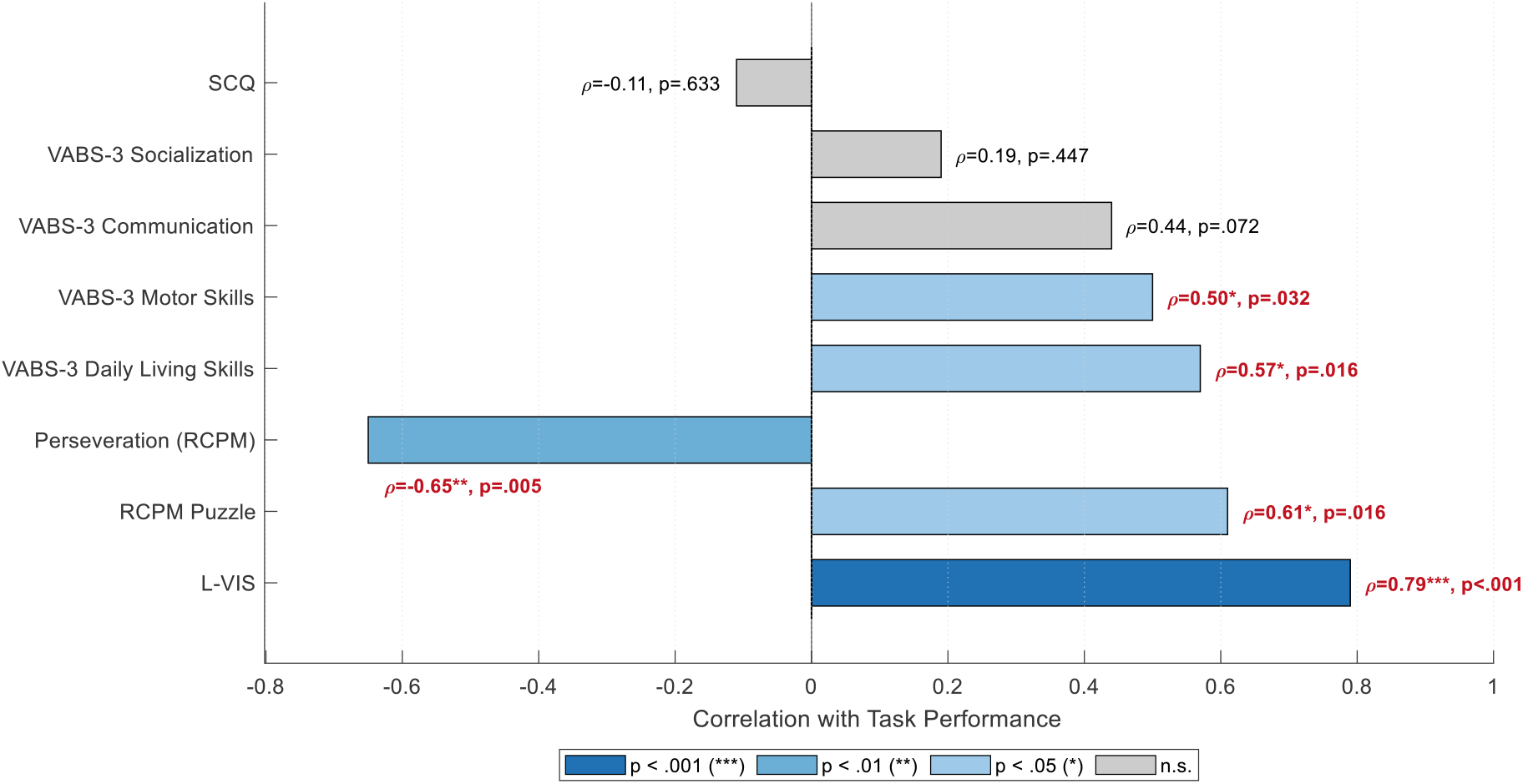
Associations between visual task accuracy and standardized measures. Bars show Spearman’s correlations with visual pointing-based task performance: RCPM, L-VIS, perseveration, SCQ, Vineland-3 domains; Socialization, Communication, Motor Skills and Daily Living Skills. Blue bars indicate significant associations, and gray bars indicate non-significant associations. P values are shown next to each bar. The p-values are FDR-corrected, as in Figure S1.

Following the model-selection rationale described in the Methods, the primary standardized-measures LME focused on communication ability (L-VIS-C) and nonverbal reasoning (RCPM). At the trial level, both predictors were positively associated with accuracy (Table 2, Table S2, Model 7): higher RCPM predicted higher accuracy (β = 0.0025, p = .037), and higher L-VIS-C scores were also associated with higher accuracy (β = 0.0478, p < .001). The fixed effects explained 19% of trial-level variance, increasing to 44% when participant- and session-level random effects were included. To assess whether this pattern reflected between-participant differences, we conducted a complementary participant-level analysis using mean accuracy per participant (Table 2, Table S2, Model 8). This analysis showed a similar pattern: L-VIS-C was a robust predictor of mean accuracy (β = 0.0506, p = .001), while RCPM was at the threshold of significance (β = 0.0026, p = .050). Together, the two measures accounted for 69% of the between-participant variance in mean accuracy.

## Discussion

This study examined visual task performance in minimally verbal autistic children (mvASD) using touchscreen oddball and contour-detection tasks. Three findings emerged. First, performance was heterogeneous, with suggested low- and high-performing subgroups. Second, performance was shaped by developmental abilities and stimulus complexity: communication ability and nonverbal reasoning accounted for approximately 69% between-participant mean-performance variance, success declined for mid-level stimuli, and RTs in HighP-mvASD largely mirrored TD patterns, unlike slower, marginally modulated RTs in LowP-mvASD. Third, exploratory findings suggested perception-action dissociations: some children fixated targets despite incorrect pointing, some successful Kanizsa performers pointed to local inducers, and more children performed within age range on nonverbal reasoning with puzzle placement than booklet pointing. Together, these findings suggest that mvASD task performance reflects not only perceptual or cognitive capacity, but also how visual information is translated into action. Unlike much of the visual-perception literature in verbal autistic individuals, the present findings show that in mvASD, apparent perceptual performance may be constrained by response format and action-selection demands.

### Methodological considerations and strengths

This study used a touchscreen paradigm that eliminated verbal instructions, allowed modelled practice with unlimited response time, and was conducted by a clinically experienced experimenter in familiar environments with individualized rewards (Kasari et al., 2013; Tager-Flusberg et al., 2017; Matsuzaki et al., 2019; Bauminger-Zviely et al., 2020). Testing across 8-10 sessions over 3-8 months allowed participants to become familiar with procedures, and high participation rates suggest that comprehension and engagement were not primary confounds. The early exclusion of six of 28 recruited children due to safety behaviors, motor limitations, vision issues, and uncertain verbal status also illustrates the practical barriers that contribute to mvASD underrepresentation in experimental and clinical research (Stedman et al., 2019; McKinney et al., 2021).

### Heterogeneity in mvASD

Children with mvASD are often treated as a single subgroup with common skill limitations (Pizzano et al., 2024; Kasari et al., 2014; Bal et al., 2016; Ben-Itzchak et al., 2014). However, prior work and the present findings challenge this assumption, showing evidence of within-age-range performance for some individuals (e.g., Pizzano et al., 2024; Courchesne et al., 2015; Bauminger-Zviely et al., 2020). In our visual perceptual tasks, high-performing mvASD participants showed reaction times comparable to TD children, and 50% of mvASD performed within age range on the RCPM puzzle board. Because subgroups were defined by accuracy, accuracy differences between them should be interpreted descriptively; however, the TD-like speed and level-sensitive RT modulation in HighP-mvASD suggest that the subgroup distinction captured a broader performance profile.

### General capacity account

Communication ability and nonverbal reasoning together accounted for approximately 69% of the between-participant variance in mean visual-task accuracy (Table 2, Model 8), supporting a general-capacity account in which broader developmental abilities contribute substantially to task success. At the trial level, however, these measures explained less variance as fixed effects alone (R^2^m = .19), although model fit improved when participant- and session-level random effects were included (R^2^c = .44; Table 2, Model 7). This indicates that trial-by-trial performance also reflected stable child-level and session-level variability not captured by the standardized measures. Thus, the general-capacity account appears substantial but incomplete. Perseveration on the RCPM puzzle correlated negatively with performance (rho =−.65, p = .005, Figure 7), suggesting that some errors may reflect response-selection or cognitive-flexibility constraints rather than limited ability or random guessing. Consistent with this interpretation, several participants who scored at floor on the pointing-based RCPM booklet, due to repeated location-based perseveration via pointing, performed within age range on the puzzle format. Similar pointing perseveration patterns were observed anecdotally during experimental tasks, warranting systematic assessment of perseveration in future studies.

### Visual complexity, saliency, and paradoxical findings

Children with mvASD showed higher accuracy for low-level than mid-level stimuli (p < .0001; e.g., 3D box and ‘Hard’ Contours, Figure 1b), consistent with reduced reliance on integrative or inference-based processing as visual complexity increases. However, this interpretation is complicated by the finding that high-performing mvASD participants reached ceiling on some mid-level tasks, including Kanizsa and light-from-above conditions. These displays may have required mid-level integration, but they may also have been solvable using local cues, such as dark/light opponency of distractors next to target in the light-from-above condition or smaller Pac-Man inducer angles in the Kanizsa condition.

A notable paradox emerged among low-performing participants in the black/white saliency task: they detected a white target more easily among gray distractors than among clearly different black distractors. Because black and white patches were both salient relative to the gray background, low-performing participants may have selected among locally salient items rather than integrating the full display to identify the oddball target. In contrast, when gray distractors appeared on a gray background, the white target was more salient and thus created a pop out effect. Better detection of circles than squares among triangular distractors further suggests sensitivity to local shape features, such as curves versus corners.

Together, these findings suggest that pointing-based performance in mvASD was strongly shaped by local saliency, contrast, and feature-level discriminability, especially when broader integration across the display was required. This aligns with saliency-map accounts, in which high local contrast guides attention (Itti & Koch, 2000), and with previous findings of atypical visual saliency and stronger attention to low-level features in autism (Wang et al., 2015).

### Perception-action dissociation

Pointing accuracy may not directly reflect perceptual ability. In the black-and-white oddball task, eye-gaze exceeded pointing among low performers, significantly for transient fixation accuracy (TFA; Figure 5a). Similar response-format effects of improved eye-gaze over pointing were observed in basic word recognition in young adults with mvASD (Ellert et al., 2025), and in nonverbal reasoning, where both the present study and Courchesne et al. (2015) showed improved performance with puzzle-board rather than pointing-based formats. Notably, although Courchesne et al. (2015) interpreted puzzle-board success as evidence of visual strengths, we suggest that the *tangible*, movable response format itself, which alters motor, spatial, and action-selection demands may be accounting for the improvement in performance.

The Kanizsa oddball task further suggests that successful detection may guide action atypically in mvASD, as nearly half of successful performers in detecting global illusory oddball pointed to local inducers (Figure 6a,b). As noted above, this local pointing could reflect local/second-order resolution of the task; given mixed findings on visual illusions in autism, this interpretation remains tentative (Hadad and Yashar, 2022). A follow-up study using the same Kanizsa stimuli and circular contours with pointing, drag-and-drop, and eye-gaze responses (Sykes-Haas and Bonneh, 2026) supports perceptual access to global structure even when spontaneous pointing suggests otherwise. Together, these findings support perception-action dissociations in mvASD, suggesting that response modality and response-selection demands can strongly affect whether perceptual or cognitive abilities are demonstrated.

### Theoretical accounts for perception-action dissociation

What could account for this hand-eye discrepancy? The following accounts are not mutually exclusive, nor are they exhaustive; rather, they are offered as hypothesis-generating to guide future investigation. These mechanisms may represent different contributors to the same broader difficulty: translating perceived targets into consistent goal-directed action. One possibility is increased motor variability. Torres et al. (2013) reported elevated stochastic variability in goal-directed and goal-less hand movements in ASD, particularly among nonverbal individuals, indicating increased motor noise. We recently extended this to oculomotor behavior, showing increased saccadic variability correlated with ADOS scores (Ziv et al., 2024). This instability, reflected in frequent short inter-saccade intervals, may lead to impulsive gaze shifts; because saccades guide hand movements (Neggers et al., 2000), such variability may cause mvASD children to miss targets that their initial transient fixations briefly located.

A second possibility concerns saliency decay and arousal. Donk et al. (2008) found that in neurotypical adults, fast saccades reliably target the most salient stimulus, whereas delayed responses show more errors, suggesting that saliency is short lived. Pupil dilation, an indicator of LC-NE system activity, reflects arousal supporting goal-directed behavior (London, 2018). Reduced pupil dilation in mvASD (Ellert et al., 2025; Bonneh et al., 2025) may therefore suggest insufficient arousal recruitment, explaining why accurate initial fixation is followed by longer reaction times and less accurate pointing.

Lidstone and Mostofsky (2021) propose that autism involves disrupted visual-motor integration (VMI), with greater reliance on proprioceptive/somatosensory feedback than visual feedback when forming internal representations for action. This account fits our gaze–pointing dissociation, but our findings refine it by suggesting that visual input is not uniformly disruptive: it can support action when targets are salient, deviant, or defined by low-level features. Despite identical motor demands across tasks, performance varied with visual characteristics, and eye-gaze showed that targets were sometimes perceived but not selected for pointing. Consistent with this, Torres et al. (2013) found that stimulus type affects stochastic variability in pointing, further suggesting that visual properties can modulate motor output in ASD. Relatedly, naturalistic grasping kinematics can reliably distinguish autistic from non-autistic adults, suggesting that visually guided action dynamics themselves carry autism-relevant information (Freud et al., 2025).

Cisek’s affordance competition hypothesis (ACH; Cisek, 2007, 2012) provides a broader framework for interpreting these findings. The model proposes that sensory information specifies multiple potential actions in parallel, which compete through mutual inhibition within fronto-parietal sensorimotor circuits and are biased by task relevance, salience, value, and contextual signals. In our oddball tasks, the possible pointing locations can be viewed as competing affordances. The gaze-pointing dissociation suggests that the target location may be detected yet not reliably selected for manual response; instead, competing distractor locations, or locally guided responses such as pointing to Kanizsa inducers, may sometimes win the competition. This dysregulation may stem from the factors discussed above, including inefficient arousal recruitment, increased motor variability, stimulus-driven attention, and impaired VMI. Together, these factors may weaken the biasing signals that normally resolve competition in favor of the task-relevant action, leading to inconsistent or atypical action selection in mvASD. This account may also relate to Robertson and Baron-Cohen’s (2017) proposal that ambiguity resolution across perceptual, language, and social-cognitive domains might rely on reciprocal inhibitory competition between neural populations. Thus, children with mvASD may ‘see the point,’ but not always point to what they see.

### Clinical implications and future directions

Despite the small sample, visual stimulus type consistently affected mvASD performance. As illustrated by the AG case in the introduction, communication may depend on how visual materials are presented. Because visual aids are widely used in mvASD intervention (e.g., Arthur-Kelly et al., 2009), future work should identify which features, saliency, deviance, or complexity, support or disrupt pointing and eye-gaze responses. This is consistent with evidence that visual stimulus properties modulate the motor output of goal-directed pointing in nonverbal autistic children (Torres et al., 2013), and that cognitive load increases corrective movements in neurotypicals (Thorpe et al., 2022), suggesting that ‘visual-perceptual’ load effects on motor output may provide an informative window into perception-action coupling in mvASD. Structured-task strengths should also not be taken to imply preserved everyday functioning. Although participants varied in reasoning, communication, and task performance, adaptive functioning remained severely impaired across the sample (Table 1).

### Limitations

Eye-tracking data were not collected from the TD reference group or high-performing mvASD participants, as performance in these groups was at or near ceiling on the black/white oddball tasks. These data would have been valuable for understanding how typical and high-performing mvASD children perform the same task through eye movements. The exploratory perception-action findings are preliminary and require replication in larger samples with dedicated eye-tracking protocols.

### Summary and conclusions

This study challenges the assumption that minimally verbal autistic children show uniformly limited visual or cognitive abilities. Although performance varied with developmental abilities and visual complexity, the findings also suggest that pointing-based responses can mischaracterize what mvASD children perceive or understand. Eye-gaze, Kanizsa pointing patterns, and puzzle-based reasoning together suggest that some children may access relevant visual or cognitive information without consistently translating it into the expected motor response. Future work should therefore examine mvASD performance not only as a measure of perceptual capacity, but as an interaction between visual input, response modality, and action-selection demands.

## Author Contribution

HSSH and YSB designed the experiments. HSSH collected the data. YSB developed the software used for running the experiment and the data analysis. HSSH and YSB analyzed the data and, HSSH wrote the manuscript, YSB revised it.

## Conflict of interest

The author(s) declared that this work was conducted in the absence of any commercial or financial relationships that could be construed as a potential conflict of interest.

## Generative AI statement

The author(s) declared that generative AI was used in the creation of this manuscript. During the preparation of this manuscript, the author(s) used Claude (Anthropic) and Chat GPT 5 to assist with language editing and formatting. After using this tool, the author(s) reviewed and edited all content as needed and take full responsibility for the content of the publication.

## Acknowledgements

We are grateful to the children and families who participated in this study, and to the staff of the specialized school for children with autism spectrum disorder for their cooperation and support. This work was supported by the Israel Science Foundation (Grant No. 657/21) and the Simons Foundation Autism Research Initiative (SFARI; Grant No. 573840), both awarded to Y.S.B.

## Appendix 1

Individual participants’ descriptive measures

Individual participants' descriptive measures
| ID mvASD | AGE | Gender | Medication | L-VIS Communication | SCQ | RCPM booklet | RCPM puzzle | shrieks | Non-verbal | Perseveration puzzle<sup>5</sup> |
| --- | --- | --- | --- | --- | --- | --- | --- | --- | --- | --- |
| Lou | 7y9m | M | N | 7 | 26 | f.p. | f.p. | N | Y | 20 |
| Lars | 8y3m | M | Y | 9 | 18 | f.p. | 38.9 | Y | Y | 8 |
| Leif | 10y8m | M | N | 2 | 28 | f.p. | 13.8 | Y | Y | 4 |
| Len | 7y5m | M | N | 8<sup>2</sup> | 26 | f.p. | f.p. | Y | Y | 19 |
| Lee | 9y5m | M | Y | 5 | 24 | f.p. | f.p. | Y | Y | 15 |
| Lara | 7y4m | F | Y | 8 | 20 | f.p. | f.p. | Y | Y | 11 |
| Liam | 10y9m | M | <sup>1</sup> | 6 | 23 | f.p. | f.p. | Y | Y | 8 |
| Lia | 6y11m | F | N | 5 | 26 | <sup>3</sup> | <sup>3</sup> | Y | Y | <sup>3</sup> |
| Leo | 9y11m | M | Y | 5 | 22 | f.p. | f.p. | Y | Y | 3 |
| Levi | 7y9m | M | Y | 9 | 27 | <sup>4</sup> | <sup>4</sup> | Y | Y | <sup>4</sup> |
| Loni | 8y3m | M | Y | 6 | 30 | f.p. | f.p. | Y | Y | 5 |
| Hal | 9y1m | M | N | 12 | 29 | 85 | 90 | N | N | 1 |
| Hila | 10y7m | F | Y | 14 | 18 | 28 | 34.4 | Y | Y | 1 |
| Haya | 10y9m | F | N | 6 | 29 | 28 | 43.8 | Y | Y | 1 |
| Hugo | 11y1m | M | Y | 12 | 20 | f.p. | f.p. | Y | Y | <sup>4</sup> |
| Hank | 6y9m | M | N | 11 | 15 | 97 | 96.7 | N | Y | 0 |
| Hugh | 8y5m | M | Y | 13<sup>2</sup> | 22 | 95 | 97.5 | Y | Y | 0 |
| Hana | 9y8m | F | Y | 8 | 23 | <sup>3</sup> | <sup>3</sup> | Y | Y | <sup>3</sup> |
| Hui | 10y4m | F | Y | 8 | 29 | f.p. | f.p. | Y | Y | 12 |
| Hans | 8y7m | M | Y | 11 | 21 | f.p. | 7.5 | Y | N | 11 |
| Holm | 8y9m | M | Y | 11 | 27 | f.p. | 27.8 | N | Y | 2 |
| Haim | 7y9m | M | Y | 9<sup>2</sup> | 25 | <sup>3</sup> | <sup>3</sup> | N | Y | <sup>3</sup> |

<sup>1</sup>Information not available <sup>2</sup>Scored by familiar staff <sup>3</sup>Unavailable for further testing <sup>4</sup>Participant unwilling to complete testing
<sup>5</sup>Perseveration: same answer location chosen on two consecutive items; RCPM never places the correct answer in the same location on adjacent items = floor performance (less than 1<sup>st</sup> percentile) = within age-range performance.

## Notes

### Competing Interest Statement

The authors have declared no competing interest.

### Summary of Updates

We are resubmitting the manuscript to correct an error in Figure 2. In the previous version, examples of the visual experimental stimuli were inadvertently omitted. This omission has now been corrected, and the revised Figure 2 has been included in the updated manuscript.

