## Supplementary materials for "Minimally verbal children with autism may ‘see the point, but do not (always) point to what they see’: A behavioral and eye-tracking study in visual perceptual processing"

#### Table S1 Visual stimulus properties

| Stimulus Properties | Color 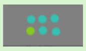 | B/W saliency 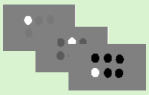 | Shapes 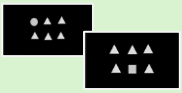 | Orientation Gratings 1st order 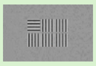 |
|---|---|---|---|---|
| Array layout | 2X3 and 2X2 | 2X3 and 2X2 | 2X3 and 2X2 | 2X3 and 2X2 |
| Stimulus Dimensions | Diameter: 3.2 cm (198px) | Diameter: 3.2 cm (198px) | Circle diameter: 2.6 cm (161px)<br>Square side: 2.6 cm (161px)<br>Triangle side :2.8 cm (173px) | Side: 3.15cm (195px) |
| Spacing * | 4.7 cm (291px) | 4.7 cm (291px) | 4.7 cm (291px) | 3.7 cm (229px) |
| Compound properties (RGB, luminance, CPD) | Oddball:<br>(0.678, 1, 0.184)<br>Distractors:<br>(0.25, 0.875, 0.182). | White: 1<br>Black: 0<br>Dark grey: 0.35<br>Mid grey: 0.47 | All stim RGB: 1 | SF**:1c/dg<br>Luminance Contrast: 32% |
| Luminance (cd/m²)<br>-Stimulus | Oddball:95<br>Distractor:90 | Oddball:145<br>Black: 3<br>Dark grey:31<br>Mid grey:61 | All stim.: 145 | All stim:73 |
| -Background | 78 | 74 | 1.5 | 110 |
| VA*** of stimulus | 3.05°-3.66° | 3.05°-3.66° | 2.67°- 3.21° | 3.01°-3.61° |
| **Stimulus Properties** | **3D BOX** 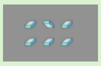 | **Shape from Shading** 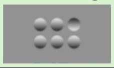 | **Kanizsa** 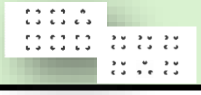 | **Orientation Gratings 2nd order** 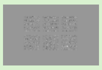 |
| Stimulus array | 2X3 | 2X3 | 2X3 | 2X3 and 2X2 |
| Stimulus Dimensions | Height:0.9 cm (56px)<br>Width:1.6cm (99px)<br>Length:1.8cm (112px) | Diameter=3.7 cm (185px) | Triangle side: 3.5 cm (216px).<br>Square side: 3 cm (185px)<br>Inducer diameter: 1.4cm (87px) | Side: 3.15cm (195px) |
| Spacing * | 4.8cm (297px) | 4.8 (297px) | 7.2cm (446px) | 3.7 cm (229px) |
| Luminance (cd/m²):<br>-Stimulus | 115 | Average:100<br>Light top:230<br>Dark bottom:10 cd/m² | Inducer:12 | SF**:1c/dg<br>Luminance contrast: 62% |
| Background | 101 | 100 | 335 cd/m² | 110 cd/m² |
| VA*** of stimulus | 2.77°-3.32° | 3.53°-4.24° | 2.86° to 3.43° | 3.01°-3.61° |

* Inter-stimulus center spacing in cm (px), non jittered. **SF= Spatial Frequency VA=Visual angle

| Properties | Contour-EASY | Contour-MEDIUM | Contour-HARD |
|---|---|---|---|
| Configurations | Low BG density* | Medium BG density* | High BG density* |
| Dimensions | Diameter:5.8 cm (313px) | Diameter:5.8 cm (313px) | Diameter:5.8 cm (313px) |
| Coordinates (x,y) | $\pm 6.8cm\ (x), 4.1cm(y)$ | $\pm 6.8cm\ (x), 4.1cm(y)$ | $\pm 6.8cm\ (x), 4.1cm(y)$ |
| Patch Contrast | Maximal | Maximal | Maximal |
| Background | 60 cd/m² | 60 cd/m² | 60 cd/m² |
| Visual angle | 5.54° to 6.64° | 5.54° to 6.64° | 5.54° to 6.64° |

*See text for details

### Table S2 linear mixed-effects models

| Model | Level | Matlab formula | Random effects | Fixed effects (β, SE, t, p) | Omnibus test | Variance explained |
|---|---|---|---|---|---|---|
| 1 | Trial | ok ~ level + (1\|obs) + (1\|obs:ses) | Intercepts: obs, ses\|obs | level: β=-0.163, SE=0.015, t(9579)=-13.66, p<.0001 | F(1,9579)=186.63, p<.0001 | — |
| 2 | Trial | ok ~ group*level + (1\|obs) + (1\|obs:ses) | Intercepts: obs, ses\|obs | group×level: β=-0.022, SE=0.030, t(9577)=-0.72, p=.47 | Not reported | — |
| 3 | Trial | RT ~ group*levelc + (1\|obs) + (1\|obs:ses) | Intercepts: obs, ses\|obs | LowP: β=235.33, SE=60.50, t=3.89, p<.001; HighP: β=16.44, p=.78; level: β=286.17, p<.0001; interactions: HighP β=-95.79, p=.02; LowP β=-270.02, p<.0001 | F(5,8664)=30.20, p<.0001 | — |
| 4 | Trial | ok ~ modality + (1+modality\|obs) | Random intercept + slope | modality: β=0.283, SE=0.102, p=.006 | F(1,218)=7.76, p=.006 | — |
| 5 | Trial | ok ~ modality + (1+modality\|obs) | Random intercept + slope | modality: β=0.192, SE=0.109, p=.08 | F(1,218)=3.10, p=.08 | — |
| 6 | Trial | amp ~ group + (1\|obs) + (1\|obs:ses) | Intercepts: obs, ses\|obs | group: β=-42.02, SE=8.89, t(1397)=-4.73, p<.0001 | F(1,1397)=22.36, p<.0001 | — |
| 7 | Trial | ok ~ RCPM + LVISC + (1\|obs) + (1\|obs:ses) | Intercepts: obs, ses\|obs | RCPM: β=0.0025, SE=0.0012, t(8078)=2.08, p=.037; LVISC: β=0.0478, SE=0.0132, t(8078)=3.62, p<.001 | — | Marginal $R^2$=.19; Conditional $R^2$=.44 |
| 8 | Subject | mean_ok ~ RCPM + LVISC | — | RCPM: β=0.0026, SE=0.0012, t(15)=2.13, p=.050; LVISC: β=0.0506, SE=0.0129, t(15)=3.90, p=.001 | — | $R^2$=.69 |

***Table S2. Full specification of linear mixed-effects models.*** *Models correspond to the LME analyses summarized in Table 2. Participant-level t-tests shown in Table 2 are not repeated here. Trial-level models included random intercepts for participant and session nested within participant unless otherwise stated.* ***Note.*** *In the MATLAB formulas, ok denotes trial accuracy coded as correct/incorrect; RT denotes reaction time; amp denotes magnitude of off-center pointing; obs denotes participant; ses denotes testing session; obs:ses denotes session nested within participant; level denotes stimulus level; levelc denotes centered stimulus level; group denotes diagnostic/performance group; modality denotes response modality (eye-gaze vs pointing); RCPM denotes Raven's Coloured Progressive Matrices score; and LVISC denotes L-VIS communication score. Random-effects terms follow MATLAB fitlme notation, such that (1|obs) specifies a random intercept for participant, (1|obs:ses) specifies a random intercept for session nested within participant, and (1+modality|obs) specifies participant-specific random intercepts and modality slopes.*

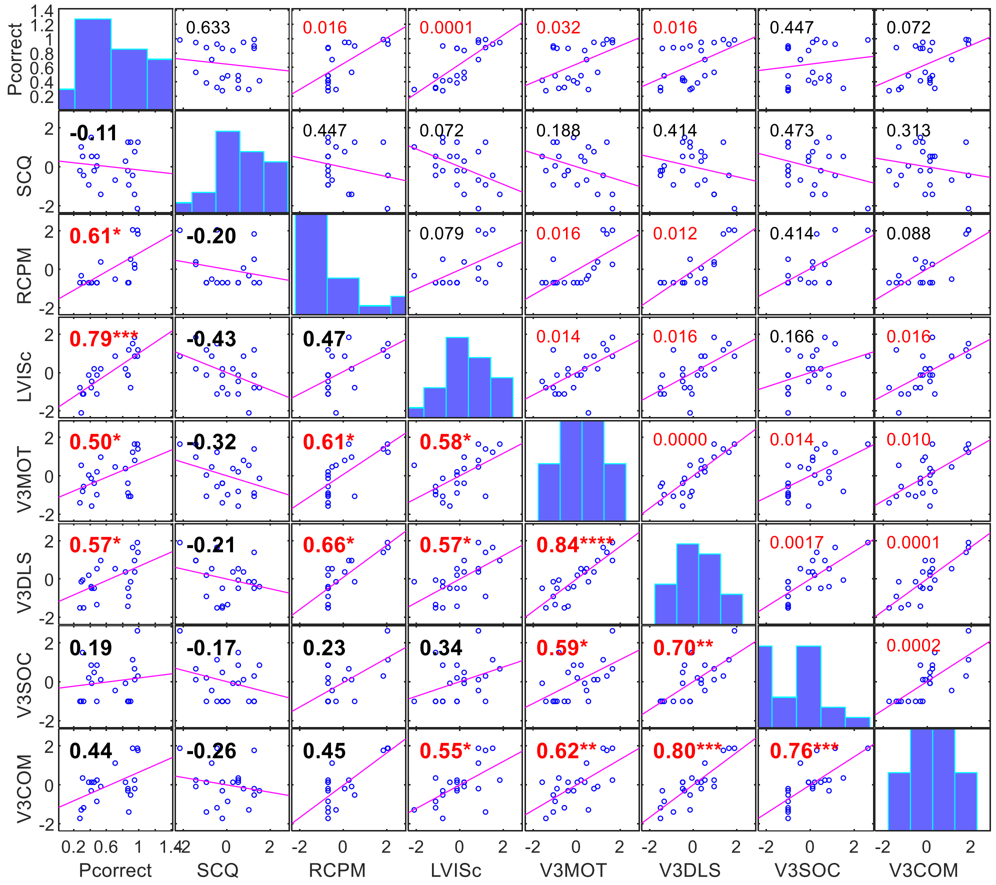


***Figure S1. Correlation matrix for visual performance and standardized developmental measures among mvASD participants.*** *Scatterplot matrix showing associations between overall visual-task accuracy (PCorrect), autism traits (SCQ), nonverbal reasoning (RCPM), communication ability (L-VISc), and Vineland-3 adaptive behavior domains: Motor Skills (V3MOT), Daily Living Skills (V3DLS), Socialization (V3SOC), and Communication (V3COM). Developmental measures are shown as z scores. Diagonal panels show variable distributions. Lower-triangle panels show Spearman correlation coefficients (ρ), and upper-triangle panels show corresponding FDR-adjusted p-values. Fitted lines are included only to visualize association direction. Red values indicate significant associations after FDR correction. Sample size was n = 22, with pairwise sample sizes ranging from 18 to 22 because of missing RCPM data.*

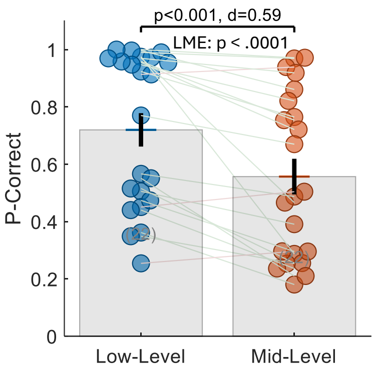


***Figure S2. Performance comparison during low and mid-level visual tasks.*** *Bee-swarm plot illustrating mvASD participants' performance in low- and mid-level visual tasks. Each pair of circles connected by a line represents one participant. The p-value from the participant-level group comparison, the LME p-value, and Cohen's d are shown above the panel.*
